# Immunomodulation of Neutrophil Granulocyte Functions by Bacterial Polyphosphates

**DOI:** 10.1101/2022.12.21.521352

**Authors:** Viola Krenzlin, Johannes Schöche, Sarah Walachowski, Christoph Reinhardt, Markus P. Radsak, Markus Bosmann

**Author notes:** Corresponding Author: Markus Bosmann, Associate Professor of Medicine, Pathology and Laboratory Medicine, Pulmonary Center, Department of Medicine, Boston University School of Medicine, Boston, Massachusetts, 02118, USA.

## Abstract

Polyphosphates are highly conserved, linear polymers of monophosphates that reside in all living cells. Bacteria produce long chains containing hundreds to thousands of phosphate units, which can interfere with host defense to infection.

Here, we report that intratracheal long-chain polyphosphate administration to C57BL/6J mice resulted in the release of proinflammatory cytokines and influx of Ly6G^+^ polymorphonuclear neutrophils in the bronchoalveolar lavage fluid causing a disruption of the physiologic endothelial-epithelial small airway barrier and histologic signs of lung injury. Polyphosphate-induced effects were attenuated after neutrophil depletion in mice. In isolated murine neutrophils, long-chain polyphosphates modulated cytokine release induced by lipopolysaccharides (LPS) from gram-negative bacteria or lipoteichoic acid from gram-positive bacteria. In addition, long-chain polyphosphates induced immune evasive effects in human neutrophils. In detail, long-chain polyphosphates down-regulated CD11b and curtailed the phagocytosis of *E. coli* particles by neutrophils. Polyphosphates modulated the migration capacity by inducing CD62L shedding resulting in CD62L^low^ and CD11b^low^ neutrophils. The release of IL-8 induced by LPS was also significantly reduced. Pharmacologic blockade of PI3K with wortmannin antagonized long-chain polyphosphate-induced effects on LPS-induced IL-8 release.

In conclusion, polyphosphates govern immunomodulation in murine and human neutrophils suggesting polyphosphates as a therapeutic target for bacterial infections to restore innate immune defense.

## Introduction

Bacterial infections cause a major burden of disease worldwide.[1] In particular, lower respiratory tract infections (i.e. pneumonia) are among the most common global causes of death.[2] Therapeutic options for life-threatening complications of these infections (e.g. acute lung injury) are limited [3], demanding better insights into the intricate relationship between pathogens and the host.

Inorganic polyphosphates are polymers of phosphate residues joined by high-energy phosphoanhydride bonds varying in length depending on the organism and tissue.[4] Polyphosphates are ubiquitous and evolutionarily conserved, exist in all living cells and serve pleiotropic functions such as energy storage, protein folding, cell proliferation, apoptosis, and coagulation.[5–7] Emerging evidence from our laboratory and others suggest that bacteria-derived polyphosphates also play a dominant role for pathogenic bacteria to evade host immune responses.[8–10] Bacterial polyphosphates are long-chains (L-PolyP) of hundreds to thousands of phosphate units, while polyphosphates released from activated mammalian platelets and mast cells are short-chains (S-PolyP) of 60-120 phosphate monomers.[7]

Neutrophil granulocytes are rapidly recruited to a site of injury and serve essential functions in innate immune defense.[11] Recent studies uncovered that platelet-type, short-chain polyphosphates mediate neutrophil-endothelial cell interactions, promote neutrophil migration, enhance recruitment and accumulation during bacterial infection, and can induce neutrophil extracellular traps.[11–16]

On the other hand, the functions of bacterial, long-chain polyphosphates on neutrophils remained undefined, and were investigated in this current report.

## Results and Discussion

### Long-chain polyphosphate-induced lung injury is neutrophil dependent

To investigate the role of polyphosphates during lung injury and inflammation, we injected long-chain polyphosphates into the trachea of C57BL/6J mice.[17] ELISA analysis of bronchoalveolar lavage fluids (BALF) 8h after the administration showed that long-chain polyphosphates induced the release of IL-6, KC, and MIP-1α (Fig. **1A**). KC is chemoattractant for several immune cells, especially neutrophils, and plays an important role in regulation of inflammatory responses. KC has a similar role as IL-8 in humans. Activated neutrophils can release MIP-1α. [18]

**Figure 1.**
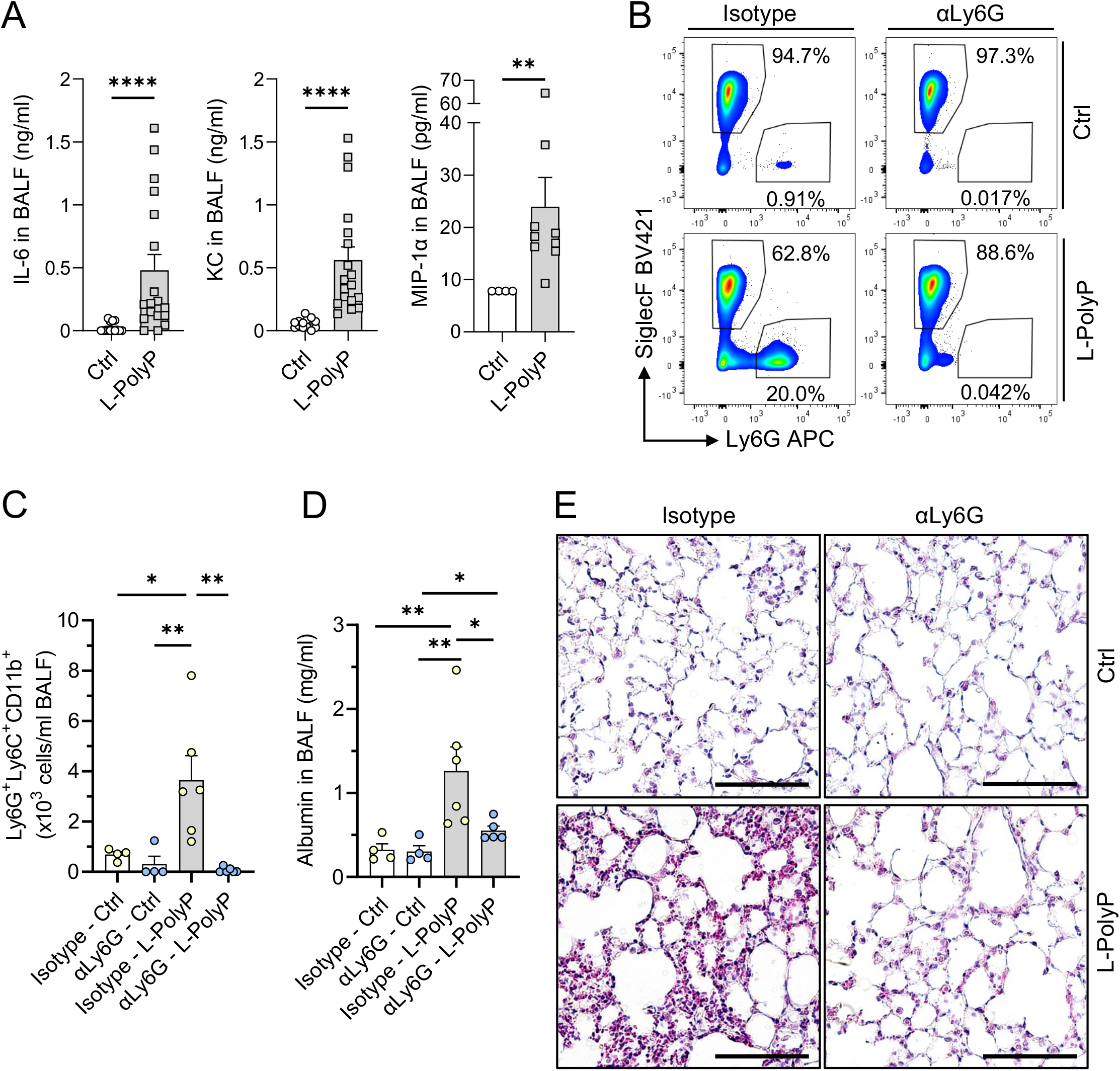
Long-chain polyphosphates cause neutrophil-dependent lung injury in mice. **(A)** IL-6, KC, and MIP-1α in broncho-alveolar lavage fluids (BALF) 8h after intra-tracheal (i.t.) administration of long-chain polyphosphates (L-PolyP) in C57BL/6J mice, ELISA. Ctrl: sham control mice. **(B-E)** Neutrophils were depleted in mice by injection of anti-Ly6G antibody (αLy6G, 100μg/mouse i.p.) 24h before the experiments. Control mice received an isotype control antibody. **(B)** Representative contour plots of Ly6G^+^ neutrophils and SiglecF^+^ cells gated from live BALF cells of isotype antibody or neutrophil-depleted mice after L-PolyP or PBS control treatment for 8h, flow cytometry. **(C)** Ly6G^+^Ly6C^+^CD11 b^+^ neutrophil count in BALF from of all mice from blots in frame B, flow cytometry. **(D)** Albumin in BALF of neutrophil-depleted and isotype-injected mice after L-PolyP or PBS for 8h, ELISA. **(E)** Representative lung sections of isotype-injected and neutrophil-depleted mice obtained 8h after L-PolyP or PBS (n=3/group); H&E staining; scale bar: 20μm. In all experiments, L-PolyP were i.t. injected in a volume of 40μl at 20mM/mouse. Sham controls (Ctrl) received PBS (40μl/mouse, i.t.). Combined or representative data from 2-3 independent experiments. Symbols represent one individual mouse (n≥4-9 mice per group) and bars indicate mean±SEM; t-test (A) or one-way ANOVA (C-D); *p<0.05; **p<0.01; ***p<0.001; ****p<0.0001.

To further study the contribution of neutrophils to polyphosphate-induced lung injury, we used a monoclonal anti-Ly6G^+^ antibody to effectively and selectively deplete neutrophils in mice before administration of long-chain polyphosphates (Supp. Fig. **1A**). Higher numbers of Ly6G^+^Ly6C^+^CD11b^+^ neutrophils (<1% vs. 20%) were present in BALF after long-chain polyphosphates (Fig. **1B**,**C**, Supp. Fig. **1B**). The polyphosphate-induced neutrophil influx caused disruption of the endothelial-epithelial lung barrier as indicated by significant differences between BALF albumin amounts of isotype-treated mice and neutrophil-depleted mice (Fig. **1D**). Histopathology of lungs showed diffuse hypercellularity caused by accumulation of neutrophils, hyaline membranes, and alveolar wall thickening consistent with pulmonary edema and acute lung injury in the long-chain polyphosphate/isotype-treated mice compared to long-chain polyphosphate/neutrophil-depleted mice (Fig. **1E**).

Thus, neutrophils were recruited by chemokines (e.g. KC, MIP-1α) and mediated polyphosphate-induced lung injury, although neutrophil influx appeared lower than for other stimuli of acute lung injury (e.g. LPS, immune complexes, C5a), where neutrophils typically comprise >80% of all BALF cells.[19, 20]

### Long-chain polyphosphates modulate LPS/LTA-induced cytokine release by murine neutrophils

To further dissect the effects of polyphosphates on mediator release, we designed experiments with isolated, purified mouse neutrophils incubated with short-chain or long-chain polyphosphates in co-stimulation with bacterial cell wall components.

First, neutrophils were incubated with LPS alone or combined with short-/long-chain polyphosphates for 7h. Long-chain polyphosphates, but not short-chain polyphosphates, significantly enhanced the LPS-induced release of several chemokines/cytokines including KC, MIP-1α, MIP-2, IL-10, IL-18, and TNFa as analyzed by multiplex assay (Fig. **2A**). In addition, separate ELISAs were performed for KC and MIP-1 a. LPS alone induced a moderate increase of these mediators which was boosted by long-chain polyphosphates at all time points studied (Fig. **2B**). As the effects were most prominent at 7h, this time point was used for further studies. RT-qPCR and ELISA confirmed the increase of the LPS-induced release of MIP-1α and KC (Fig. **2C, D**). The co-presence of long-chain polyphosphates significantly increased the gene expression of these chemokines, while no consistent effects for short-chain polyphosphates were observed (Fig. **2C, D**). Similar results were noted for lipoteichoic acid (LTA), a major constituent of the cell walls of gram-positive bacteria, although LTA was less potent compared to LPS (Fig. **2C, D**). In conclusion, bacterial-type, long-chain polyphosphates alone were without effects, but primed murine neutrophils to amplify cytokine release, when bacterial pathogen-associated molecular patterns were present as co-stimulus.

**Figure 2.**
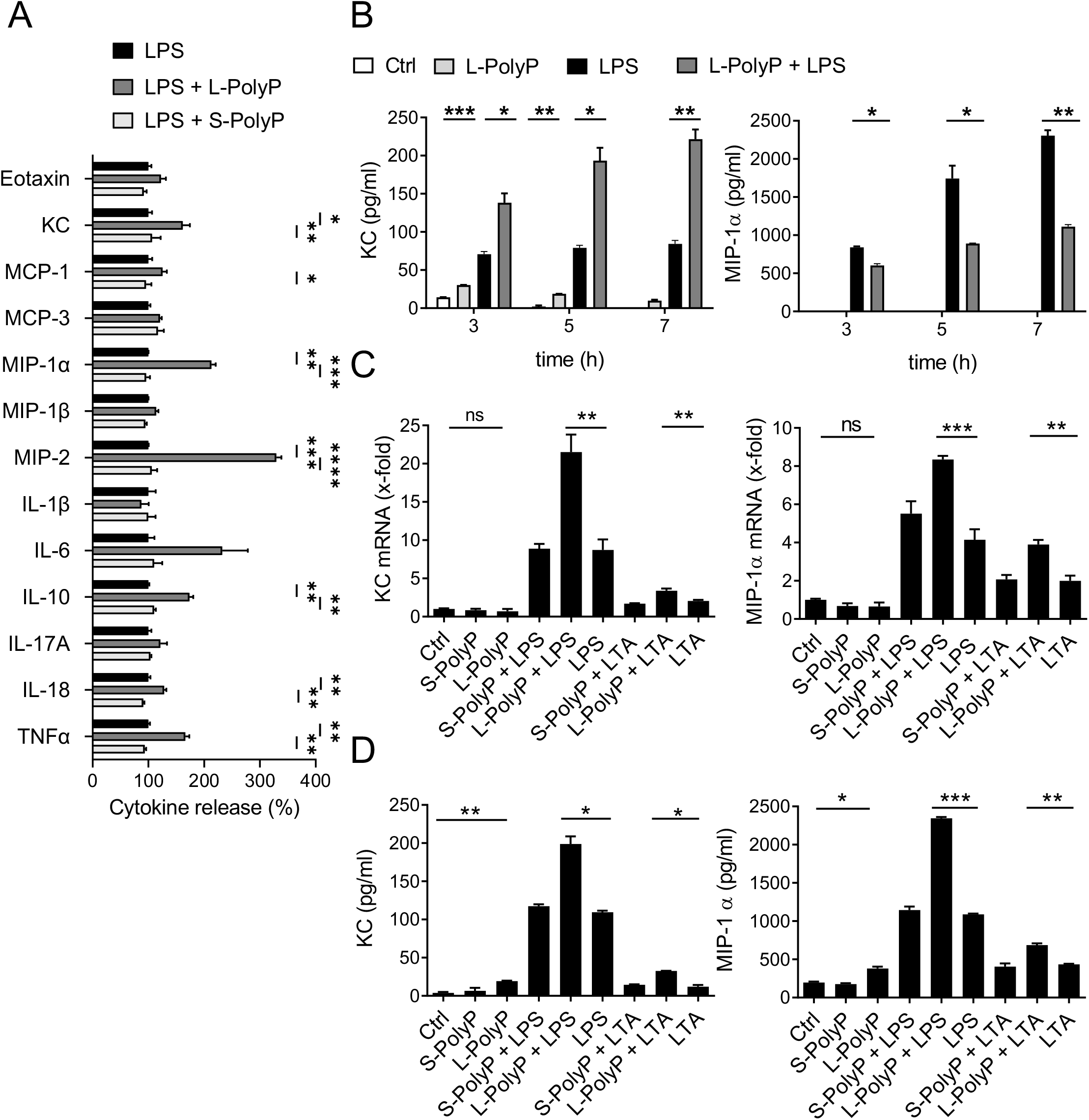
Long-chain polyphosphates increase LPS-induced cytokine release by murine neutrophils. **(A)** Multiplexed bead-based assay of supernatants from isolated peritoneal neutrophils (C57BL/6J) after incubation with LPS ± L-/S-PolyP for 7h. The release of each cytokine after LPS alone was set as 100% value. Data are pooled from 2 independent experiments. **(B)** Isolated neutrophils were incubated with LPS ± L-/S-PolyP for indicated time periods before MIP-1α and KC ELISA. Resting neutrophils served as controls (Ctrl). **(C-D)** Isolated neutrophils were incubated with LPS or LTA ± L-/S-PolyP for 7h. (C) RT-qPCR and (D) ELISA of MIP-1α and KC. The concentrations for all experiments were: L-PolyP (50μM), S-PolyP (50μM), LPS (100ng/ml), LTA (10μg/ml). B-D are representative of n≥3 independent experiments (each n>3). Data are presented as mean±SEM; two-way ANOVA (A) or t-test (B-D), *p<0.05; **p<0.01; ***p<0.001; ****p<0.0001; ns: not significant.

### Long-chain polyphosphates mediate immune evasive effects in human neutrophils

Next, we sought to investigate if long-chain polyphosphates also act on human neutrophils. In addition to the production of inflammatory mediators, the host defense functions of neutrophils include the regulation of activation markers, degranulation and phagocytosis.[21] The activation markers CD11b (integrin αM, subunit of CR3), CD62L (L-Selectin) and CD66b were measured by flow cytometry on isolated human neutrophils stimulated for 1h with short-/long-chain polyphosphates ± LPS/LTA. Long-chain polyphosphates induced down-regulation of CD11b from the neutrophil surface (Fig. **3A**). This was not observed with LPS or LTA alone (Fig. **3A**). In contrast, LPS and LTA alone induced robust shedding of CD62L, which was even more pronounced after co-incubation of LTA and long-chain polyphosphates (Fig. **3B**). While long-chain polyphosphates alone induced significant shedding of CD62L and increased CD66b, short-chain polyphosphates alone did not have any effect on CD62L/CD66b surface expression (Fig. **3B**). In general, CD11b^high^CD62L^low^ neutrophils are the phenotype considered activated for firm vascular adhesion and extravasation towards a chemotactic gradient.[22] While CD62L downregulation suggests that long-chain polyphosphates rapidly activate neutrophils, the CD11b^low^ phenotype limits their transepithelial migration capabilities. CD66b and CD11b are stored in different neutrophil granules, which allows their reciprocal regulation.[21]

**Figure 3.**
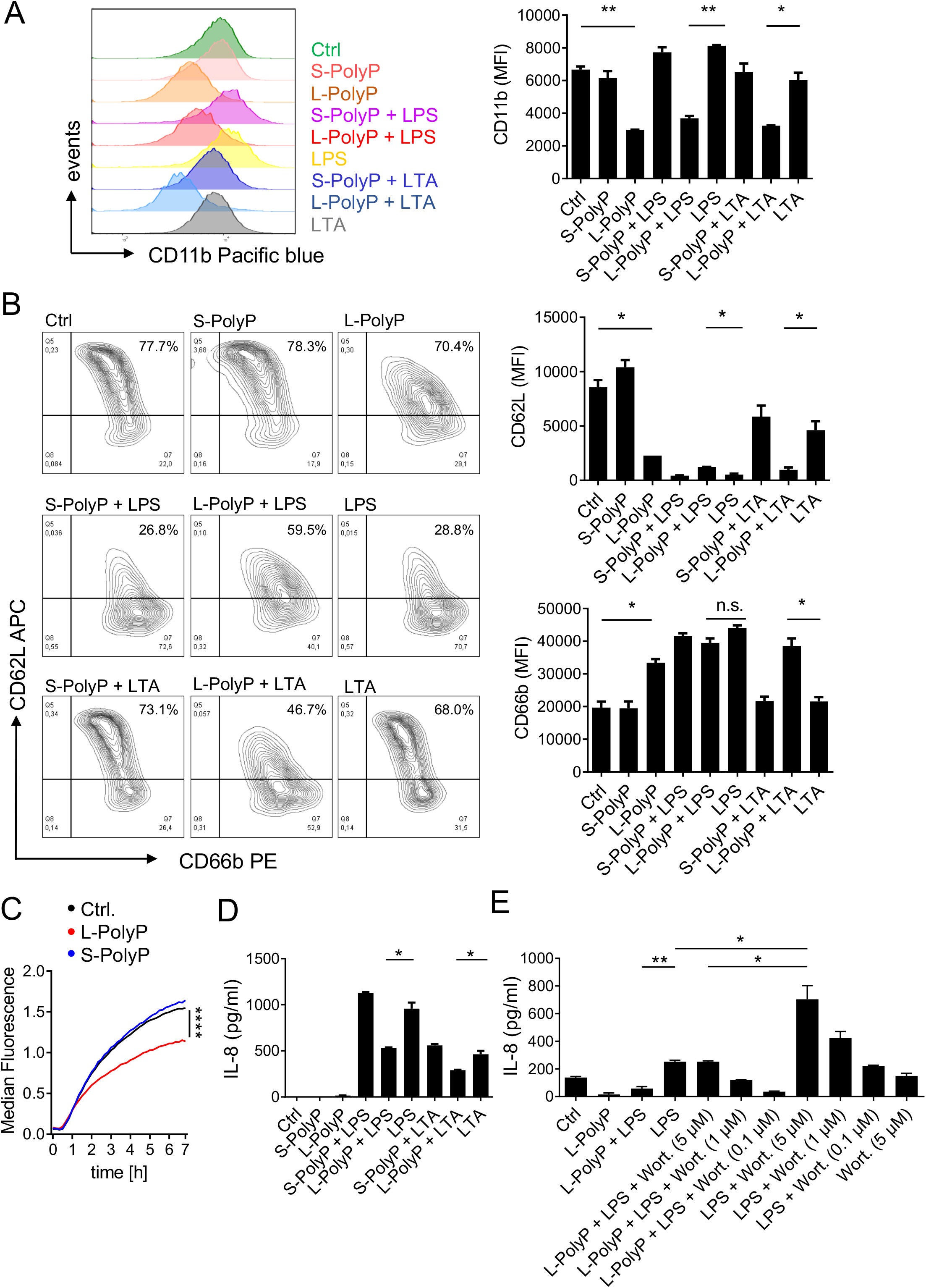
Long-chain polyphosphates mediate immune evasive effects in human neutrophils. **(A-B)** Isolated human neutrophils were stimulated for 1h with LPS, LTA, L-PolyP, S-PolyP alone or in combination. Surface expression of CD11b, CD62L, and CD66b are presented as representative histograms/plots and mean fluorescence intensities (MFI), flow cytometry. **(C)** Phagocytosis was evaluated by incubation of human neutrophils with pHrodo *E. coli* particles (333μg/ml) ± L-/S-PolyP and fluorescence was measured over time. **(D)** IL-8 from human neutrophils stimulated for 7h with LPS, LTA, L-/S-PolyP as indicated, ELISA. **(E)** IL-8 from human neutrophils stimulated with LPS, LTA, L-/S-PolyP and Wortmannin (Wort.) as indicated, 7h, ELISA. Concentrations for all experiments were: L-PolyP (50μM), S-PolyP (50μM), LPS (100ng/ml), LTA (50μg/ml) Wort. (5, 1 0.1 μM). All data are representative of n≥3 independent experiments (each n>3) and presented as mean±SEM; t-test (A,B,D,E) or one-way ANOVA (C),*p<0.05; **p<0.01, ****p<0.0001; ns: not significant.

Since CD11b also serves functions in phagocytosis, isolated human neutrophils were studied in a phagocytosis assay using conditionally fluorescent pHrodo *E. coli* particles ± short-/long-chain polyphosphates. Interestingly, long-chain, but not short-chain polyphosphates significantly reduced the uptake of *E. coli* by human neutrophils (Fig. **3C**).

### Long-chain polyphosphates attenuate LPS-induced IL-8 secretion via the PI3K/Akt signaling pathway

Binding of LPS to TLR4 on neutrophils activates the PI3K/Akt pathway, which limits release of IL-8 and other cytokines. PI3K/Akt signaling is also induced by polyphosphates.[17, 23] Here, long-chain polyphosphates alone were not sufficient to promote the IL-8 release by human neutrophils (Fig. **3D**). In contrast, LPS-induced IL-8 production was significantly and selectively inhibited by long-chain polyphosphates (Fig. **3D**). This implied that important signals transmitted by IL-8 are weakened, such as chemotaxis of other neutrophils or monocytes, and the autocrine/paracrine activation of phagocytes. Similarly, long-chain polyphosphates also antagonized the LTA-induced IL-8 release (Fig. **3D**). To investigate the relevant signaling pathway, PI3K was blocked by the small molecule inhibitor, wortmannin (Fig. **3E**). Wortmannin antagonized long-chain polyphosphate-dependent suppression of LPS-induced IL-8, indicating that long-chain polyphosphate control IL-8 release via the PI3K/Akt signaling pathway (Fig. **3E**). Wortmannin dose-dependently enhanced IL-8 release after LPS alone, confirming that PI3K/Akt is an inhibitory pathway in terms of IL-8 production (Fig. **3E**).

### Concluding Remarks

In conclusion, we demonstrate that polyphosphates, in chain-lengths that predominantly exist in bacteria, modulate critical host defense functions of mouse and human neutrophils. Despite their primitive structure, the effects of polyphosphates are versatile and distinct for each endpoint studied. Targeting polyphosphates could restore natural immunity against infections and prevent their life-threating complications, acute respiratory distress syndrome and sepsis.[9]

## Material and methods

### Ethics approval statement for human and animal studies

The local ethics committee (Landesaerztekammer Rheinland-Pfalz, Mainz, Germany) approved this study (No. 837.258.17 (11092)) in accordance with the Declaration of Helsinki. Deidentified blood was obtained from male healthy volunteers (22-40 years) who gave their informed consent.

All studies with mice were approved by the State Investigation Office of Rhineland-Palatinate and the Institutional Animal Care and Use Committee (IACUC) of Boston University. C57BL/6J mice (8-12-week-old) were housed in a specific pathogen-free environment. Sex-age matched cohorts were used for experiments.

#### Polyphosphate-induced lung injury

Mice were anesthetized with ketamine/xylazine and polyphosphates slowly injected intratracheally (i.t.) as described before.[17] BALF was collected in 1ml ice-cold PBS supplemented with 2mM EDTA (Gibco, Life Technologies, Grand Island, USA) and 1X proteases inhibitor (EASYpack Complete Mini EDTA-free cocktail tablet, Roche, Wilmington, USA). After centrifugation, BALF cells were stained for flow cytometry and cell free-supernatants were cryopreserved (−80°C) before further analysis.

#### Neutrophil depletion

Mice received a single i.p. injection of 100μg anti-Ly6G antibody (αLy6G, clone 1A8, BioXCell, Lebanon, USA) or isotype control antibody (clone 2A3, BioXCell) in 200μl PBS, 24h before polyphosphates.

#### Histology

Lungs were perfused with PBS and inflated with 500μl freshly prepared 4% paraformaldehyde (#18505, Ted Pella Inc, Altadena, USA) for fixation, and paraffin-embedded sections were stained with hematoxylin and eosin (H&E). Images were captured on a Leica DM4 B light microscope with integrated LAS X software (Leica Microsystems GmbH, Wetzlar, Germany).

#### Isolation and purification of murine peritoneal neutrophils

Mice were i.p. injected twice (−20h/−3h) with 1ml of 9% (w/v) casein in sterile saline before euthanasia.[24] The peritoneal lavage fluids (5ml HBSS) were collected in a sterile syringe, filtered through a 100μm filter, and centrifuged for 5min, 1500rpm at 4°C. The supernatants were removed, and cells diluted in MACS buffer (Miltenyi Biotec, Bergisch Gladbach, Germany) to 1×10^8^/ml and Ly6G^+^ cells were isolated according to the anti-Ly6G MicroBead Kit (#130-092-332, Miltenyi Biotec). The cells were labeled with biotinylated anti-Ly6G antibodies (anti-Ly6G-biotin, 1:1000), mixed with anti-biotin microbeads (1:40) and separated over a MACS column. The magnetically retained Ly6G^+^ cells were eluted as the positively selected cell fraction (purity: ≥98% Ly6G^+^ cells by flow cytometry; viability: ≥95% by trypan blue).

#### Isolation and purification of human neutrophils

EDTA blood (9ml) was drawn from peripheral veins of healthy volunteers and subjected to the MACSxpress Whole Blood Neutrophil Isolation Kit (#130-104-434, Miltenyi Biotec). The blood was incubated with labeling antibodies and placed in the magnetic separator. The labeled cells adhered to the tube wall and erythrocytes sedimented. The supernatant containing the undisturbed neutrophils was transferred to a new tube, and erythrocyte depletion cocktail (#130-098-196, Miltenyi Biotec, 1:50) was added. Finally, the tube was again placed in the magnetic separator for 15 min and the undisturbed target cells (purity: ≥97% CD66b^+^ cell by flow cytometry; viability: ≥95% by trypan blue) were recovered in the flow through.

#### Detection of proteins

Mouse MIP-1α/CCL3 (#DY450), mouse KC (#DY453-05), mouse IL-6 (#DY406), human IL-8 (#DY208-05) (R&D Systems, Minneapolis, USA) and mouse albumin (#E99-134, Bethyl Laboratory, Waltham, USA) were quantified by ELISA, and other mediators by 22-plex ProcartaPlex mouse basic kit (Thermo Fisher Scientific, Waltham, USA) on a Luminex-200 instrument (BioRad, Hercules, USA), all according to manufacturer’s instructions.

#### Real-time qPCR

RNA isolation and qPCR were performed as reported [17]. Primer sequences: mouse MIP-1α: (forward) 5’-CCAAGTCTTCTCAGCGCCAT-3’, (reverse) 5’-TCCGGCTGTAGGAGAAGCAG-3’; mouse KC: (forward) 5’-TCTCCGTTACTTGGGGACAC-3’, (reverse) 5’-CCACACTCAAGAATGGTCGC-3’; mouse GAPDH: (forwardv) 5’-TACCCCCAATGTGTCCGTCGTG-3’, (reverse) 5’-CCTTCAGTGGGCCCTCAGATGC-3’.

#### Flow cytometry

The cells were washed in FACS buffer (1% BSA, 0.01% sodium azide, 2.5mM EDTA in sterile PBS; 650g, 10min, 4°C) and preincubated for 15min with anti-CD16/CD32 Fc-block antibody (10μg/ml; BioLegend, San Diego, CA, 1:50) in FACS buffer followed by a 30min incubation with fluorescence dye-conjugated, anti-mouse antibodies for SiglecF (clone E50-2440, 1:400), CD11c (clone N418, 1:100), CD11b (clone M1/70, 1:100), Ly6G (clone 1A8, 1:50, clone 1A8, 1:400), Ly6C (clone HK1.4, 1:80/1:600), F4/80 (clone BM8, 1:100), CD62L (clone DREG-56, 1:80), CD66b (clone G10F5, 1:40) and corresponding isotype controls (all from BioLegend). Analysis was done in adherence to guidelines [25].

#### Phagocytosis assays

Isolated human neutrophils (1.5×10^5^ cells/100μl) were challenged with 333μg/ml pHrodo™ green-E. coli (Thermo Fisher Scientific) in the presence or absence of polyphosphates at 37°C in the Fluoraskan Ascent FL fluorometer (Thermo Fisher Scientific). The fluorescence signals were acquired every 5min for 7h.

#### Reagents

Polyphosphates were a gift from Dr. James H. Morrissey and Dr. Stephanie A. Smith.[9, 17, 23] LPS from *E. coli* (O111:B4, Sigma-Aldrich, St. Louis, USA), LTA (Sigma-Aldrich), and Wortmannin (Cell Signaling Technology, Danvers, MA).

#### Statistical analysis

The GraphPad Prism Version 9 software was used for statistical analysis. All values are expressed as mean and error bars represent SEM. Data sets comparing two groups were analyzed by two-tailed Student’s t-test and multiple comparisons were made with ANOVA. In vitro experiments were performed independently 2-3 times and are shown as representative or pooled data. *In vivo* experiments were done with the numbers of mice per group as indicated by symbols (circles, squares, diamonds) in the figures. Differences were considered significant when p<0.05.

## Abbreviations

L-PolyP: long-chain polyphosphates
LPS: lipopolysaccharides
LTA: lipoteichoic acid
BALF: bronchoalveolar lavage fluids

## Data sharing and data availability

The data that support the findings of this study are available from the corresponding author upon reasonable request.

## Author contributions

V.K, J.S., and S.W. designed and performed experiments, and analyzed data. M.P.R. provided guidance for neutrophil isolation protocols and helpful comments. C.R. provided comments. M.B. and V.K. wrote the manuscript, which was edited by S.W., M.P.R. and C.R. M.B. conceived and supervised the study, designed experiments, interpreted data, and provided funding.

## Acknowledgements and funding statement

This work was supported by financial resources obtained from the Federal Ministry of Education and Research (01EO1503 to M.B.), the Deutsche Forschungsgemeinschaft (BO3482/3-3, BO3482/4-1 to M.B.; CRC156 TP KS01, CRC1066 TP B18, CRC1292/2 Project No. 318346496 TP21N to M.P.R.), and the National Institutes of Health (R01AI153613 to M.B.). C.R. acknowledges funding from the Forschungsinitiative Rheinland-Pfalz and ReALity. C.R. was awarded a fellowship of the Gutenberg Research College at the Johannes Gutenberg-University Mainz. C.R. is a DZHK scientist and a principal investigator in the BMBF Cluster4Future CurATime. We thank Foruzandeh Samangan, Melissa Pesta, Anne Hinds and Markus Dudek for technical assistance. We thank Kara Farquharson for reading the final version of the manuscript. The authors are responsible for the content of this publication.

