## Supporting_information for "Immunomodulation of Neutrophil Granulocyte Functions by Bacterial Polyphosphates"

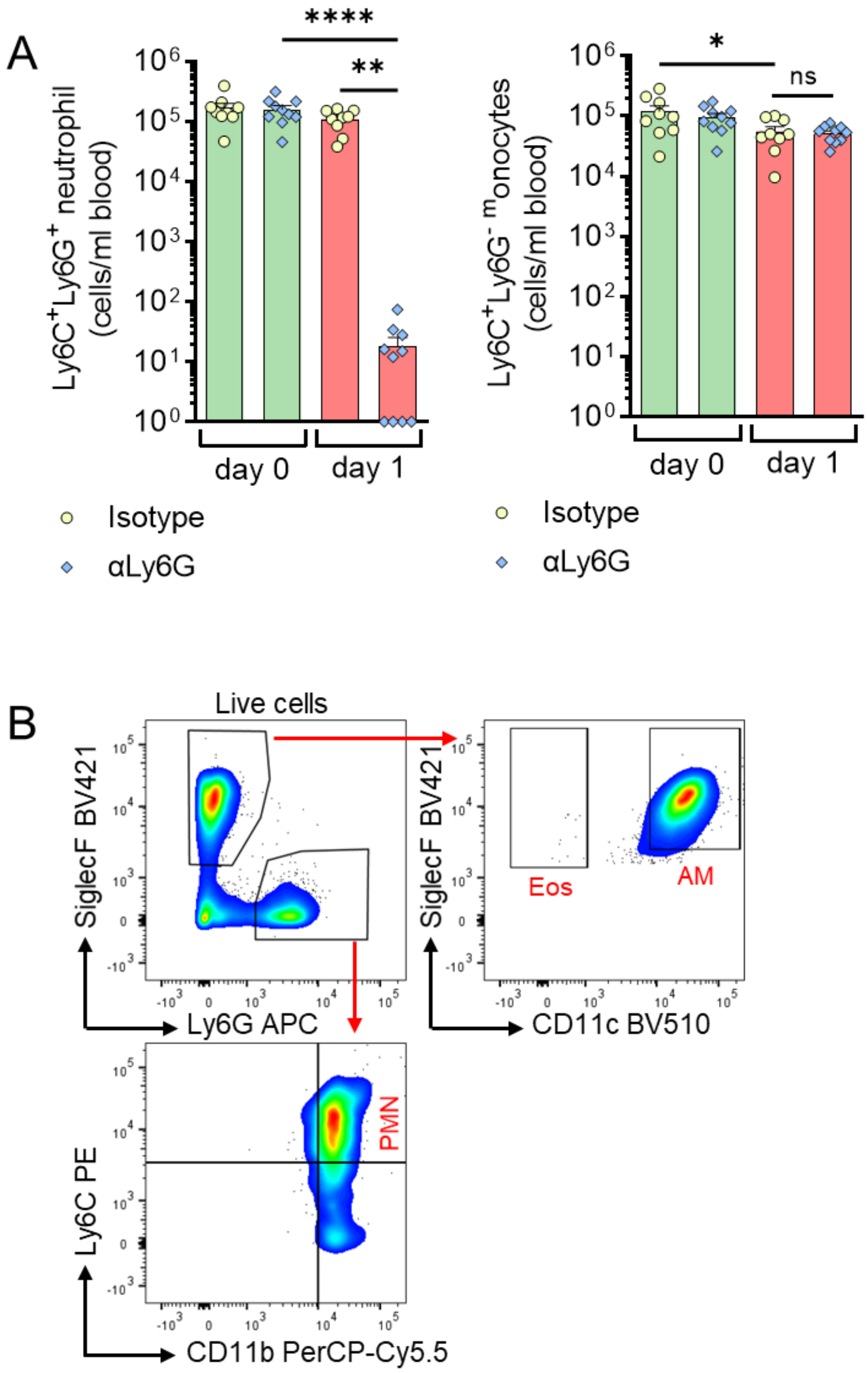


**Supplemental Figure 1. Flow cytometry gating strategy and specific depletion of neutrophils. (A)** $Ly6C^+Ly6G^+$ neutrophils and $Ly6C^+Ly6G^-$ monocytes in blood of wild type mice before (day 0) and 24 h (day 1) after i.p. administration of depleting anti-Ly6G antibody (αLy6G, 100 µg/mouse) or isotype matched control antibody, flow cytometry. **(B)** Gating strategy for BALF immune cells following intratracheal (i.t.) administration of L-PolyP (40µl at 20mM) in C57BL/6J mice. Representative plots by flow cytometry of alveolar macrophages (AM), neutrophils (PMN) and eosinophils (Eos) identified 8h after instillation are shown. Data are representative of 2 independent experiments and are shown as mean ± SEM values from the analysis of n≥4-9 mice per group; one-way ANOVA; *p<0.05; **p<0.01; ****p<0.0001, ns: not significant.
